# Effects of focused ultrasound in a “clean” mouse model of ultrasonic neuromodulation

**DOI:** 10.1101/2023.05.22.541780

**Authors:** Hongsun Guo, Hossein Salahshoor, Di Wu, Sangjin Yoo, Tomokazu Sato, Doris Y. Tsao, Mikhail G. Shapiro

## Abstract

Recent studies on ultrasonic neuromodulation (UNM) in rodents have shown that focused ultrasound (FUS) can activate peripheral auditory pathways, leading to off-target and brain-wide excitation, which obscures the direct activation of the target area by FUS. To address this issue, we developed a new mouse model, the double transgenic Pou4f3+/DTR × Thy1-GCaMP6s, which allows for inducible deafening using diphtheria toxin and minimizes off-target effects of UNM while allowing effects on neural activity to be visualized with fluorescent calcium imaging. Using this model, we found that the auditory confounds caused by FUS can be significantly reduced or eliminated within a certain pressure range. At higher pressures, FUS can result in focal fluorescence dips at the target, elicit non-auditory sensory confounds, and damage tissue, leading to spreading depolarization. Under the acoustic conditions we tested, we did not observe direct calcium responses in the mouse cortex. Our findings provide a cleaner animal model for UNM and sonogenetics research, establish a parameter range within which off-target effects are confidently avoided, and reveal the non-auditory side effects of higher-pressure stimulation.

## INTRODUCTION

Focused ultrasound (FUS) has the potential to modulate cortical and deep-brain regions with a spatial resolution on the scale of millimeters, considerably more precise than established non-invasive neuromodulation technologies such as transcranial magnetic stimulation and transcranial current stimulation.^1–19^ However, FUS can activate peripheral auditory pathways and cause off-target activation throughout the brain, including both ipsilateral and contralateral regions, regardless of the specific brain targets being stimulated.^20–24^ These auditory confounds have become a constant challenge in ultrasonic neuromodulation (UNM) experiments in rodents. To eliminate these confounds, using deaf animal models is an effective approach. However, surgical or chemical methods of deafening can be invasive or cause systemic toxicity, which limits the potential for fully awake and long-term chronic experiments.^20,21,25^ A recent UNM study used genetically deaf knockout mice with deficits in the inner hair cells,^26^ but their congenital deafness may impair brain plasticity and cortical development, potentially limiting their utility in experiments involving sensory processing, learning, and other cognitive functions.^27,28^ Therefore, it remains challenging to non-surgically deafen adult normal-hearing mice without compromising their capability for neural recording and behavioral experiments.

Here we address this challenge by developing a new “clean” mouse model for ultrasound (US) research and characterizing the safe parameter range for future UNM and sonogenetics studies. This double transgenic mouse model (Pou4f3^+/DTR^ × Thy1-GCaMP6s) allows us to use diphtheria toxin (DT) to non-invasively ablate all the hair cells without causing systemic toxicity to the mice. Additionally, by expressing the calcium indicator GCaMP6s in the brain, we can simultaneously read neuronal activity across different cortical regions in awake, deafened mice during UNM.

Using this new mouse model, we show that FUS-elicited auditory confounds and off-target brain activation can be effectively reduced or eliminated up to certain pressures in fully awake, deafened mice. As the pressure is increased, the fluorescence signal at the FUS focus gradually decreased due to focal heating and the fluorophore’s thermal dependence. Additionally, high-pressure FUS elicited enhanced visual responses to external stimuli (LED flashes) or extensive contralateral brain activation, possibly due to the activation of non-auditory peripheral sensory receptors. When the pressure exceeded 1600 kPa, we observed strong calcium responses originating from the FUS focus and subsequently propagating throughout the ipsilateral hemisphere. This strong depolarization was associated with brain damage, as confirmed by histology, indicating that high-pressure FUS causes damage-evoked seizure-like spreading depolarization. These findings provide key insights for future UNM and sonogenetics studies. First, they offer an effective deaf mouse model where the sensory confounds of UNM can be eliminated up to a certain pressure. Second, they delineate a FUS parameter space where non-auditory confounds and tissue damage can be minimized. Third, with thermal fluorescence dimming, they provide a convenient approach to visualize the focus of FUS in the cortex.

## RESULTS

### Diphtheria toxin deafens double transgenic Pou4f3^+/DTR^ x Thy1-GCaMP6s mice

Transgenic Pou4f3^+/DTR^ (Pou) mice were bred with Thy1-GCaMP6s (Thy1) mice to produce double transgenic Pou4f3^+/DTR^ × Thy1-GCaMP6s (PouThy1) mice, expressing the heterozygous human diphtheria toxin receptor from the endogenous Pou4f3 locus,^29^ and the fluorescent calcium indicator GCaMP6s.^30^ The PouThy1 mice maintain normal hearing and balance until being treated with DT (**Figure 1A**), which ablates all their inner and outer hair cells.^29^ The deafness of the treated mice can be examined by imaging cortical auditory responses through a clear skull (**Figures 1B and 1C**). In awake mice, we compared the auditory responses to audible broadband noise and visual responses to light flashes among three groups, which were Thy1 mice treated with DT (Thy1-DT), PouThy1 mice treated with saline (PouThy1-saline), and PouThy1 mice treated with DT (PouThy1-DT). Audible sound (90 dB SPL) activated the auditory cortex and other cortical regions in Thy1-DT and PouThy1-saline groups while no such activation pattern was observed in the PouThy1-DT group (**Figures 1D-1F**), suggesting DT can effectively deafen PouThy1 mice. In contrast, light flashes (80 ms in duration) evoked similar calcium responses in the visual cortex among all the three groups, suggesting that DT does not damage non-auditory cortex (**Figure 1G**).

**Figure 1.**
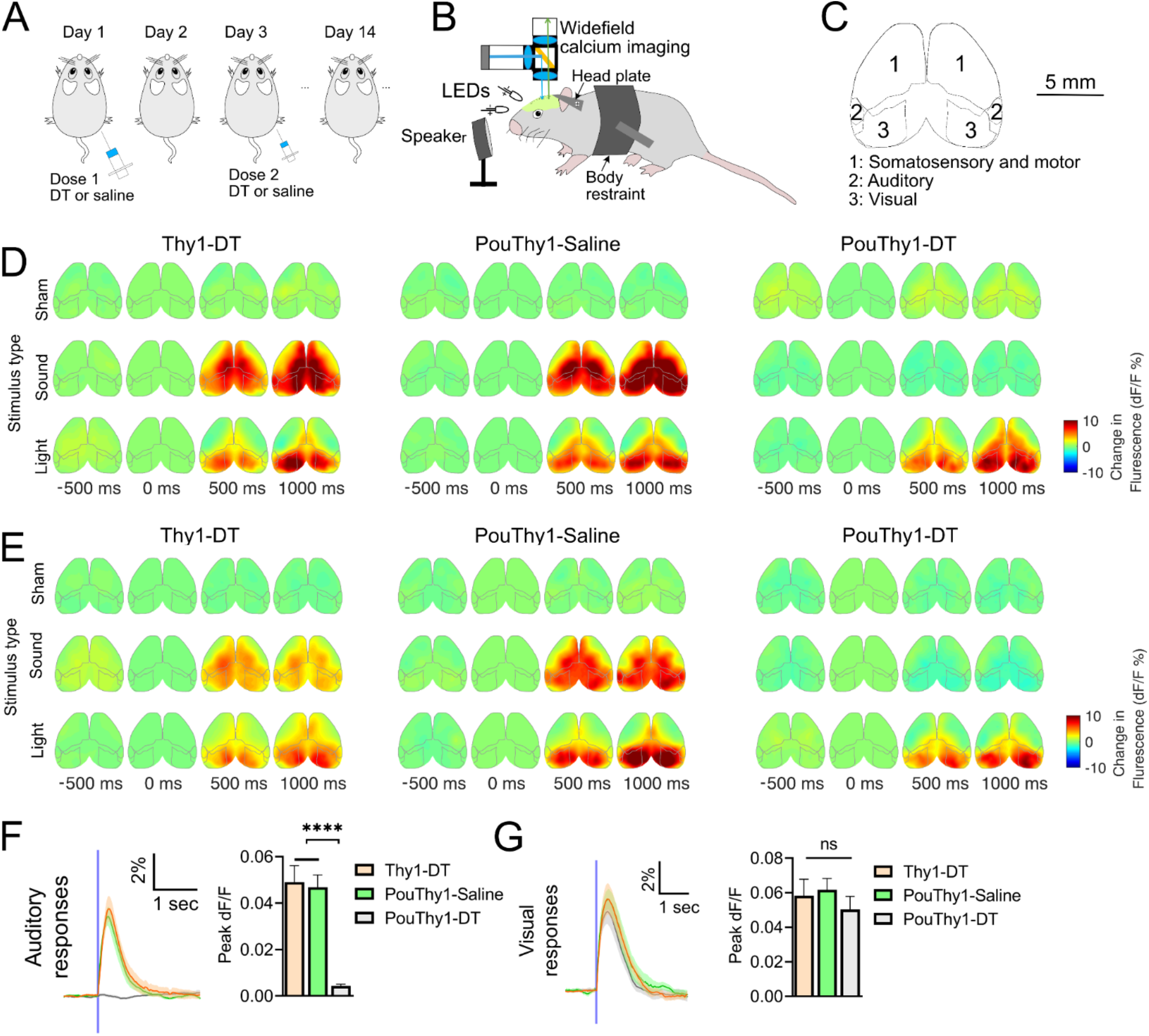
Cortical responses to sham, audible sounds, and light flashes. (**A**) Diagram of protocol for mouse deafening. Two injections of diphtheria toxin (DT) or saline were spaced two days apart. Wide-field imaging experiments were performed at least two weeks after the first injection to wait for the ablation of hair cells in the cochlear and utricles. (**B**) Illustration of wide-field calcium imaging setup. A speaker was positioned in the front of the mouse. Two LEDs were positioned to the right and left eyes, respectively. (**C**) Illustration of the top view of the cortex from Allen Mouse Brain Common Coordinate Framework (CCFv3). The visual, auditory, somatosensory, and motor cortices were indexed with numbers. (**D** and **E**) Representative examples of cortical activation map to sham, light flashes to both eyes, audible broadband noises to both ears. Two animals are presented for each group. (**F**) Auditory responses to audible sounds and the peak dF/F of the Thy1-DT, PouThy1-saline, and PouThy1-DT groups (n = 10 mice for each group, one-way ANOVA ****p < 0.0001, Tukey’s post comparison). (**G**) Visual responses to light flashes and peak dF/F of visual responses of the Thy1-DT, PouThy1-saline, and PouThy1-DT groups (n = 10 mice for each group, one-way ANOVA p = 0.1050, ns is not significant). Mean trace is solid and SEM is shaded. Bar graph values represent mean ± SEM.

### Deafening reduces or eliminates off-target widespread cortical activation to ultrasound

We further investigated if the PouThy1-DT mouse model could eliminate the auditory confounds and the off-target widespread cortical activation reported by previous UNM studies.^20,21^ Our in vivo UNM setup comprised a wide-field camera and a single element transducer angled to the brain for simultaneous FUS stimulation and calcium imaging.^21^ The tip of the transducer (fundamental frequency at 270 kHz and third harmonic at 916 kHz) was manually aligned to the stimulation target, which is -0.5 mm anterior and 2.5 mm lateral (**Figure 2A**). We tested a continuous FUS waveform with a pulse duration (PD) of 500 ms, which was shown to effectively activate cultured cortical neurons via specific mechanosensitive calcium channels.^31^ Additionally, similar waveforms have also been used in various animal and human UNM experiments.^3,20,22,24,26,32–36^

**Figure 2.**
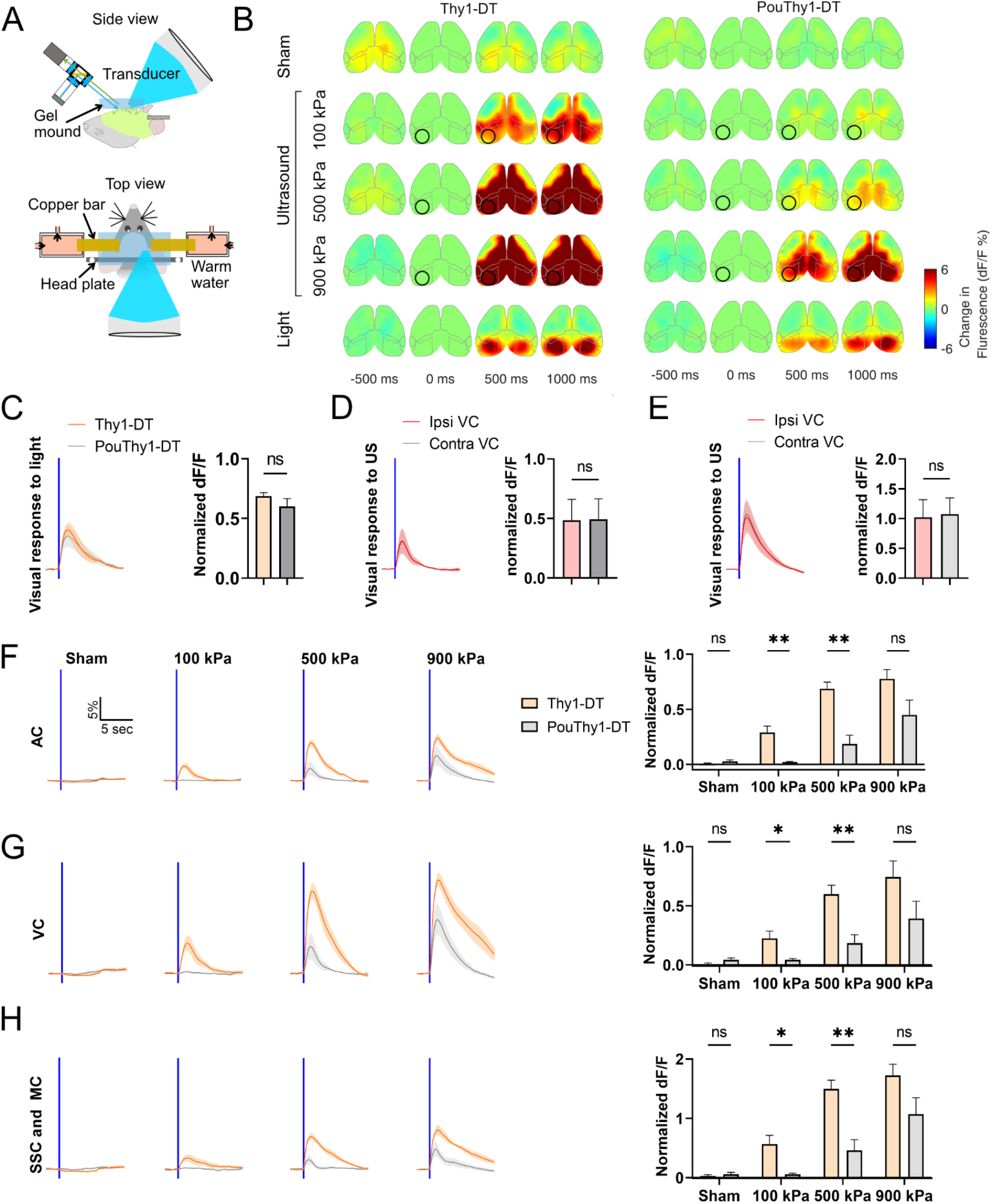
Deafening reduces off-target responses to 270 kHz ultrasound. (**A**) Illustration of simultaneous ultrasonic neuromodulation (UNM) and wide-field cortical imaging. The ultrasound (US) transducer and the imaging equipment were both angled at 45 degrees from parallel to allow optical access to the focus. The transducer was immersed in a cone filled with degassed water. The cone was then coupled to the skull with degassed US gel, which was flattened with a glass plate. The temperature of the gel mound was regulated at approximately 35 °C using bilateral copper bars via heat conduction. The other ends of the bars were sealed in 3D printed tubes and submerged in circulating warm water to maintain a constant temperature. (**B**) Representative examples of cortical activation map from one normal hearing mouse (Thy1-DT) and one deafened mouse (PouDTR-DT) to sham, US (270 kHz center frequency, 500 ms PD, pressure at 100, 500, and 900 kPa), and light flashes. The US target zone is shown as a black circle. The boundary maps are the same as in Figure 1. (**C**) Visual responses to light flashes of normal hearing and deafened mice, and the normalized peak dF/F of the two groups (n = 6 mice for each group, unpaired t-test, two-tailed, p = 0.2656). (**D** and **E**) Cortical responses at ipsilateral (ipsi) US focus and its contralateral (contra) counterpart to US at 500 kPa (D) and 900 kPa (E) and light flashes. The normalized peak dF/F are compared (n = 6 animals, unpaired t-test, two-tailed, p = 0.9755 and 0.8999 for D and E, respectively). (**F**-**H**) Auditory, visual, somatosensory and motor responses to sham and US at different pressures. Normalized peak dF/F are compared between the two groups (n = 6 animals for each group, unpaired t-test, two-tailed, *p < 0.05, **p < 0.01, ns is not significant). Mean trace is solid and SEM is shaded. Bar graph values represent mean ± SEM.

As expected, when using the 270 kHz FUS, we were able to observe strong auditory confounds and off-target brain activation in normal hearing mice (Thy1-DT). In contrast, in deafened mice (PouThy1-DT), the widespread cortical responses were mostly eliminated when using low peak negative pressure FUS (≤ 500 kPa). Unexpectedly, we still clearly observed bilateral off-target brain activation from non-auditory regions when increasing the pressure to 900 kPa (**Figures 2B and 2F-2H**). To examine if this activation was directly elicited by FUS, we compared the ipsilateral and contralateral responses of the visual cortex to FUS, but we did not find that the ipsilateral focus had stronger responses than its contralateral counterpart (**Figures 2D and 2E**). This suggests that the activation is not likely to be caused directly by local FUS stimulation of the targeted visual cortex. Among possible indirect mechanisms, since DT administration completely ablates hair cells in our transgenic model,^37^ the observed off-target brain excitation may be caused by the activation of peripheral non-auditory receptors such as cutaneous receptors and/or photoreceptors by the relatively wide beam of the 270 kHz FUS (∼8.85 mm lateral full width at half maximum [FWHM] for pressure) as these receptors are excitable to FUS stimulation.^34,38,39,41^

To assess the impact of focal zone size, we stimulated with the third harmonic of our transducer, at 916 kHz, with 1.4 mm FWHM laterally for pressure so that the focus can be confined within one hemisphere. As expected, in normal hearing mice, the application of FUS to the visual cortex resulted in widespread cortical activation at pressure as low as 100 kPa. In contrast, in fully deafened mice, the off-target brain activation was eliminated at pressures up to 900 kPa (**Figures 3 and S2**). Taken together, these results reinforce the importance of using fully deafened mice for UNM experiments because FUS can produce strong auditory and widespread brain activation in normal hearing mice at pressure as low as 100 kPa. In addition, they reveal the additional concern of non-auditory side-effects when using low-frequency transducers with large focal zones relative to brain size.

**Figure 3.**
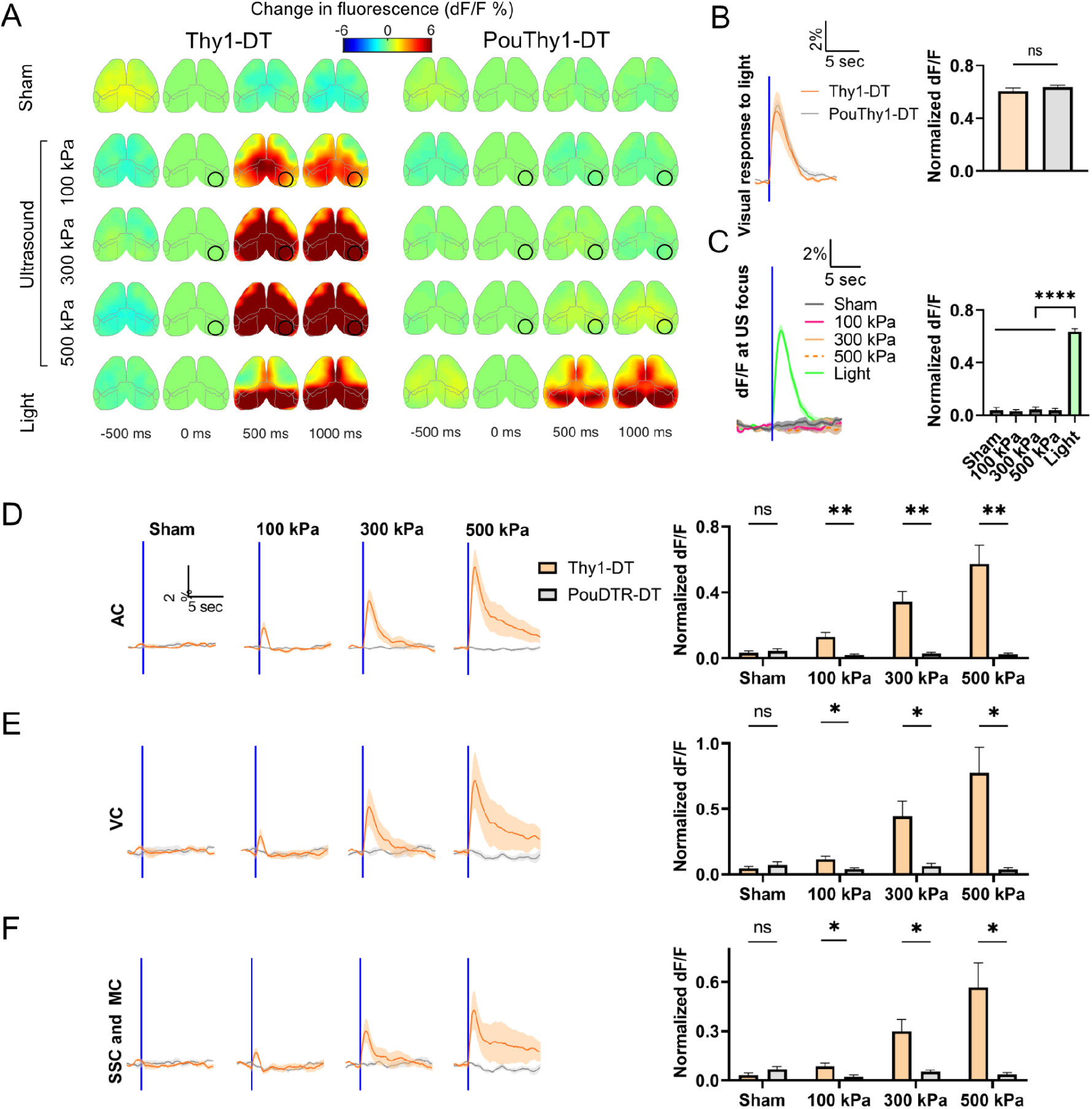
Deafening eliminates off-target responses to 916kHz ultrasound. (**A**) Representative examples of cortical activation map from one normal hearing mouse (Thy1-DT) and one deafened mouse (PouDTR-DT) to sham, US (916 kHz, 500 ms PD, pressure at 100, 300, and 500 kPa), and light flashes. The US target zone is shown as a black circle. (**B**) Visual responses to light flashes of normal hearing and deafened mice, and the normalized peak dF/F of the two groups (n = 6 animals for each group, unpaired t-test, two-tailed, p = 0.8618). (**C**) Cortical responses at US focus to sham, US at different pressures, and light flashes, and normalized peak dF/F (n = 6 animals, one-way ANOVA ****p < 0.0001, Tukey’s post comparison). (**D**-**F**) Auditory, visual, and somatosensory and motor responses to sham and US at different pressures. Normalized peak dF/F are compared between the two groups (n = 6 mice for each group, unpaired t-test, two-tailed, *p < 0.05, **p < 0.01, ns is not significant). Mean trace is solid and SEM is shaded. Bar graph values represent mean ± SEM.

### Ultrasound reduces focal fluorescence in deafened mice via a thermal mechanism

In the clear skull preparation, we noticed a decrease in local calcium indicator fluorescence following high pressure (900 kPa, 500 ms PD) FUS stimulation (**Figure S2**), which is in line with the temperature-dependent fluorescence dips reported in another recent UNM study.^33^ To better characterize the relationship between continuous FUS heating reporter fluorescence without the potentially distorting effects of the skull, we replaced bilateral skull regions with TPX, an acoustically and optically transparent polymethylpentene plastic, 0.125 mm in thickness.^40^ The windows covered +3 to - 4 mm AP, +1 to +5 mm ML for each hemisphere (**Figure 4A**). The large windows can accommodate the full 916 kHz FUS focus and allow imaging of major portions of the bilateral motor, somatosensory, and visual cortices.

**Figure 4.**
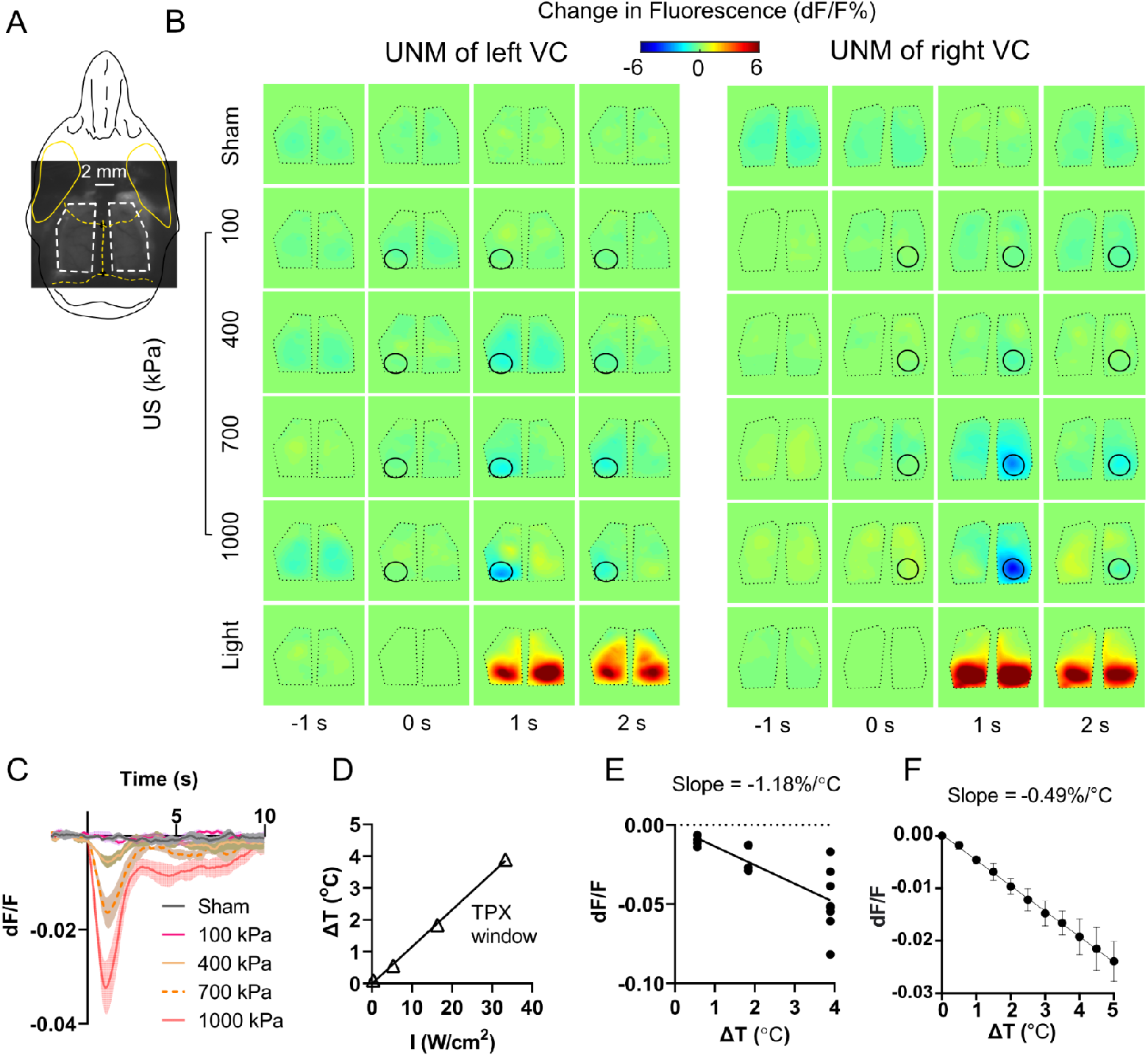
Temperature dependence of fluorescence in in vivo deafened mouse brain and in vitro human cells. (**A**-**B**) Two representative examples of cortical activation map in response to sham, US at different pressures (916 kHz, 500 ms PD, pressure at 100, 400, 700, 1000 kPa), and light flashes. A raw image of a TPX window mouse is shown in A. The length and posterior width of each window are approximately 7 and 4 mm, respectively, allowing for complete acoustic access to the brain for the 916kHz US. The US target zone is shown as a black circle. The write/black polygonal dots represent the boundaries of the TPX windows. (**C**) Focal calcium responses to sham and US at different pressures (i.e., 100, 400, 700, 1000 kPa) across four animals. Mean trace is solid and SEM is shaded. (**D**) In vivo measurement of the brain temperature increases during UNM. Intensities of 0.33, 5.33, 16.33, 33.33 W/cm^2^ correspond to pressures of 100, 400, 700, 1000 kPa, respectively. The measured temperature increase linearly correlates with the US intensity (R^2^ = 0.999). (**E**) Fluorescence change is plotted against measured temperature increase with a slope of –1.18%/°C for in vivo brain. (**F**) Fluorescence change is plotted against measured temperature increase with a slope of -0.49% for in vitro HEK293T cells.

When increasing the peak positive pressure from 100 kPa to 1000 kPa in increments of 300 kPa, we observed the decrease of focal fluorescence following the onset of FUS, which then gradually returned to the baseline (**Figures 4B and 4C**). We measured the temperature change at the brain surface under the TPX window with an optical fiber sensor during UNM in a separate animal (**Figure 4D**) and calculated our in vivo preparation having a temperature dependence of fluorescence of approximately –1.18%/°C (**Figure 4E**), relatively consistent with Estrada’s result of −1.9 ± 0.7%/°C in brain slices.^33^ To study the effects of temperature on cellular GCaMP fluorescence, we heated a population of GCaMP6f-expressing human HEK293 cells using a qPCR machine from 37 °C to 42 °C in 1-degree increments and simultaneously measured cellular fluorescence. We found a linear decrease in fluorescence intensity when the temperature increased (**Figure 4F**), suggesting that the observed ultrasound-induced fluorescence dimming in vivo could, at least in part, be attributed to the ultrasound-induced elevation in temperature.

### Ultrasound elicits long-lasting negative fluorescence signals at the focus and bilateral off-target brain activation in deafened mice

As we observed long-lasting fluorescence dimming following the onset of FUS at 1000 kPa (**Figure 4C**), we further increased the interstimulus interval (ISI) to 150 s to investigate the recovery time (defined in Methods) of the focal fluorescence signal at 1000 kPa and 1300 kPa. We found that the recovery time for FUS at 1000 kPa was 10.45 s, similar to that when the ISI was 20 s. In contrast, the recovery time for 1300 kPa was 69.6 s (**Figure 5C**), much longer than the time required for the brain temperature to return to the baseline, which was 55.36 s (**Figure S5**). Moreover, in some animals, we observed off-target ipsilateral decrease of fluorescence (example 1 in **Figure 5B**). These findings cannot be explained by thermal confounds, and may therefore indicate that some neuronal inhibition may also take place, as reported in other studies.^33,35^

**Figure 5.**
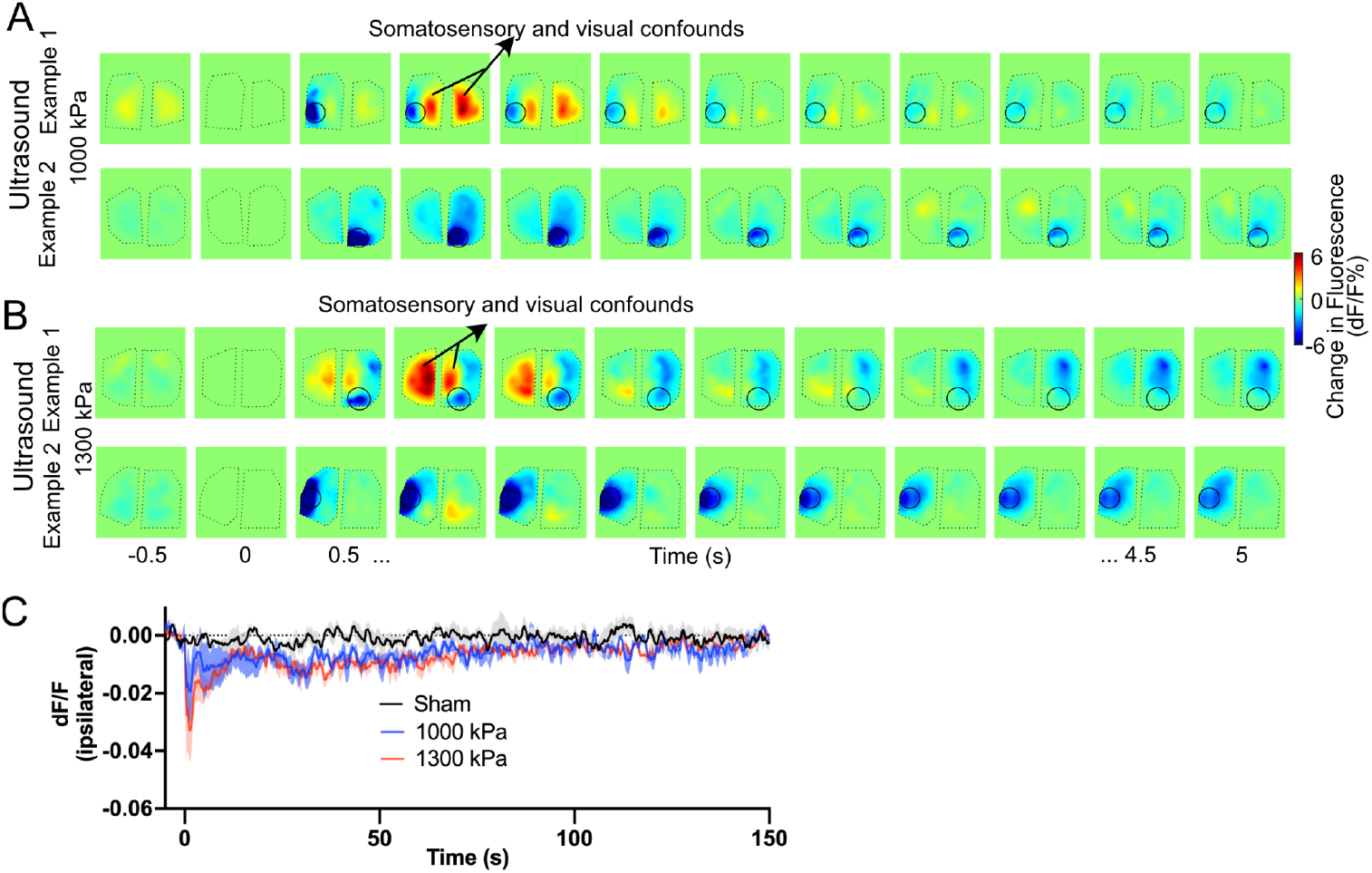
Off-target cortical responses and localized fluorescence decrease to ultrasound in deafened mice. (**A**-**B**) Representative examples of cortical activation map in response to US (916 kHz, 500 ms PD) at 1000 kPa (A) and 1300 kPa (B), respectively. For each pressure, one example (with black arrows) shows bilateral off-target brain activation in conjunction with localized decrease of fluorescence signal (one out of four animals for each group) while the other one shows only the decrease of the fluorescence signal at the focus. The off-target brain activation may be due to activation of ascending somatosensory and visual pathways. A complex cortical activation pattern, including focal and ipsiteral off-target fluorescence dimming and bilateral off-target brain activation, can be observed. The US target zone is shown as a black circle. The black polygonal dots represent the boundaries of the TPX windows. (**C**) Focal ipsilateral calcium responses slowly returned to the baseline over a period of approximately 100 seconds (n=4 animals for each group). The interstimulus interval in these experiments was 150 s.

Example 2 in Figures 5A and B demonstrates a general response pattern of the cortical activation map in most animals, where FUS elicited strong ipsilateral decrease of fluorescence signals due to heating and potential neuronal inhibition. However, interestingly, in a few animals, we observed a more intricate response pattern, where the focal fluorescence dimming coincided with bilateral off-target brain activation,^34,39,41^ which were elicited by FUS heating and potential activation of ascending visual and somatosensory pathways, respectively. The dimming partially countered the ipsilateral activation, thereby making it appear weaker than contralateral the side (example 1 in **Figures 5A and 5B**). A similar cortical activation pattern can also be found when performing paired FUS and visual (light flash) stimulation, where FUS activated bilateral peripheral visual pathways, boosting the contralateral visual-evoked calcium responses (VERs) and countering the ipsilateral dimming caused by FUS heating (**Figure S3**).

### High pressure ultrasound elicits focal depolarization

Having failed to obtain evidence of direct brain activation in awake deafened mice with FUS at pressures up to 1300 kPa, we further tested if pressures of 1600 kPa and above (in increments of 400 kPa at 916 kHz) could elicit stronger calcium responses, potentially overcoming the negative fluorescence dip caused by heating. At this higher pressure, we observed a very strong calcium signal at the focus (∼200% peak dF/F, **Figure 6**), which was ∼ 20-fold larger than that elicited by sensory stimuli (e.g., **Figure 1**). Originating at the FUS focus, this excitation propagated throughout the ipsilateral hemisphere over approximately one minute (**Figure 6**). This spatiotemporal pattern is similar to that elicited by focal cortical seizures.^42,43^ On the contralateral side, there was a relatively weaker (about 15% peak dF/F) brain activation having similar duration with the ipsilateral response, which could be explained by activation of peripheral somatosensory and visual pathways and/or by propagating ipsilateral activation across the corpus callosum, which will warrant further investigation. We also found that the strong focal calcium signal was not readily repeatable in the same animal. As shown in an example (**Figure 6C**), when performing five consecutive FUS trials (Stim 1 to 5) at an interval of 10 minutes in one experiment, only the first and third trials resulted in signal, indicating potential tissue damage. To determine effects on the tissue, we perfused three animals 24 hours after they received a single pulse of US stimulation (2000 kPa, 500 ms PD, 916 kHz) and performed H&E staining, with two animals showing clear damage in the area of stimulation (**Figure 6E**). The combined live imaging and histological evidence suggests that the strong propagating calcium signal was due to focal neuronal damage leading to spreading activation over the ipsilateral cortex, rather than direct non-damaging US stimulation.

**Figure 6.**
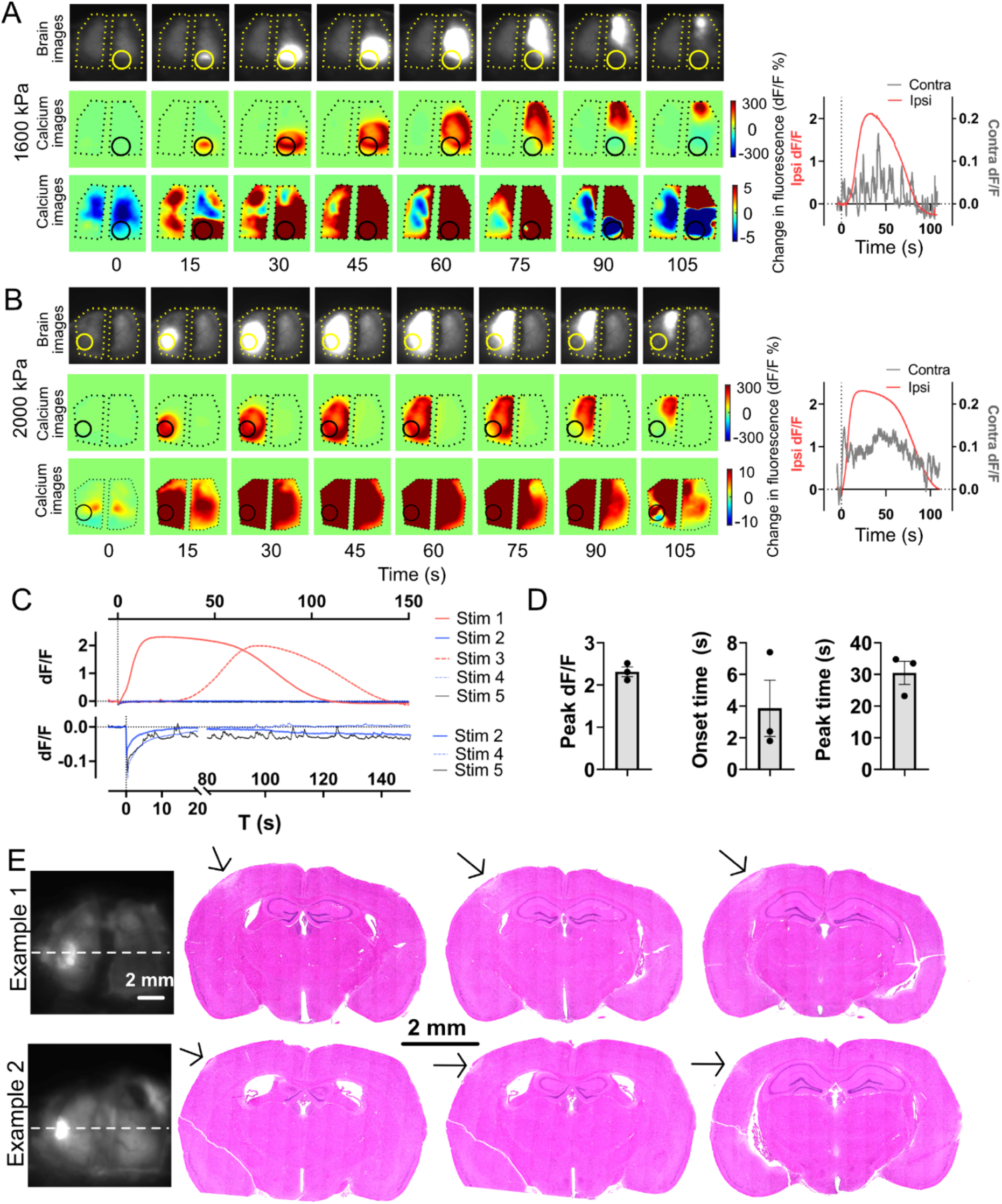
Ultrasound produces localized and hemispherically spreading disruptive depolarization. (**A** and **B**) Two representative examples of cortical activation maps of localized depolarization by US in deafened animals. The depolarization started at the focus and then propagated over the ipsilateral hemisphere. The right panels are the dF/F of the ipsilateral focus (red curve) and its contralateral counterpart (black curve). The ipsilateral dF/F is approximately 20 folds of the contralateral dF/F. The US target zone is shown as a yellow/black circle. The black polygonal dots represent the boundaries of the TPX windows. (**C**) The dF/F at the focus in response to five consecutive US stimulation (2000 kPa, 500 ms PD) with an interval of 10 min. The zoomed curves are shown below. Only trial #1 and #3 elicited depolarization. (**D**) The peak dF/F is 2.31±0.20, onset time is 30.5 ± 6.35 s, and peak time is 3.87 ± 3.07 s, respectively (n =3 animals). Bar graph values represent mean ± SEM. (**E**) Examples of H&E staining at the sonication site suggest that high-pressure US (2000 kPa, 500 ms PD) induced brain damage (white regions pointed by black arrows) at the focus.

### Low-pressure pulsed ultrasound does not elicit localized calcium signals

After comprehensively characterizing responses to continuous FUS, we also tested pulsed stimulation parameters (**Table 1**) informed by UNM studies that have reported that low-intensity pulsed FUS can elicit electrophysiological activity at the FUS focus^2,44^. To keep the FUS focus on the ipsilateral target and minimize somatosensory effects, we used 916 kHz FUS. We did not observe any activation in our deafened mice (**Figure 7**), even with 3-4 fold higher pressure than used in previous reports. In contrast, we observed strong bilateral brain activation in normal hearing mice using the same parameters (**Figure S4**), suggesting that auditory confounds may contribute to the widespread brain activation elicited by pulsed FUS.

**Table 1.** Pulsed ultrasound parameters investigated

| Parameter |  | f (MHz) | c/p | PD (ms) | PRF (Hz) | Duty cycle | Np | US Stim length (ms) | Pr (kPa) | I<sub>SPPA</sub> (mW/cm<sup>2</sup>) | I<sub>SPTA</sub> (mW/cm<sup>2</sup>) |
| --- | --- | --- | --- | --- | --- | --- | --- | --- | --- | --- | --- |
| Set1 | US1 | 0.916 | 210 | 0.229 | 2500 | 57.3% | 150 | 60 | 100 | 330 | 189 |
|  | US2 | 0.916 | 210 | 0.229 | 2500 | 57.3% | 150 | 60 | 300 | 3000 | 1719 |
|  | US3 | 0.916 | 183 | 0.2 | 1500 | 30% | 120 | 80 | 100 | 330 | 99 |
|  | US4 | 0.916 | 183 | 0.2 | 1500 | 30% | 120 | 80 | 300 | 3000 | 900 |
| Set2 | US5 | 0.916 | 18 | 20 | 30 | 60% | 2 | 67 | 72.6 | 175.80 | 105.48 |
|  | US6 | 0.916 | 18 | 20 | 30 | 60% | 2 | 67 | 300 | 3000 | 1800 |
|  | US7 | 0.916 | 2 | 2 | 300 | 60% | 20 | 67 | 72.6 | 175.80 | 105.48 |
|  | US8 | 0.916 | 2 | 2 | 300 | 60% | 20 | 67 | 300 | 3000 | 1800 |

**Figure 7.**
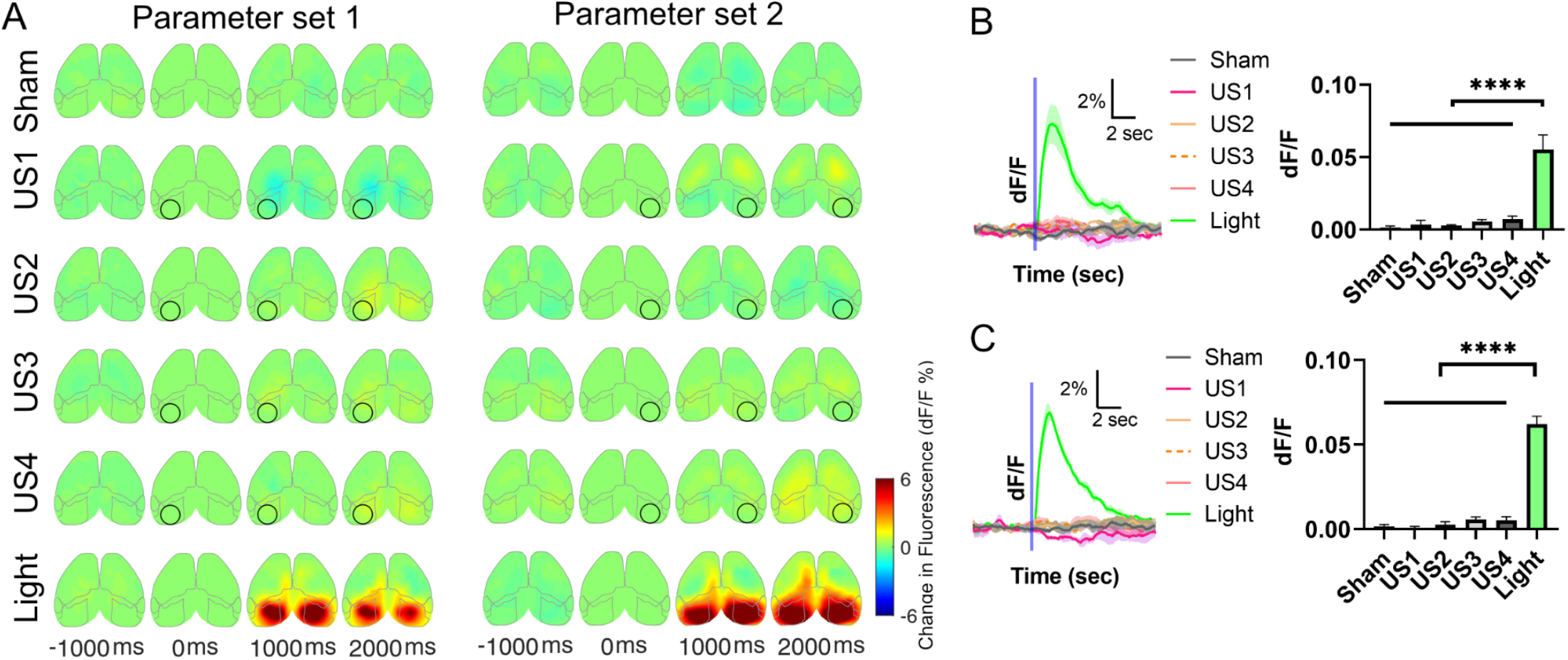
Cortical responses to low-pressure pulsed ultrasound. (**A**) Two representative cortical activation maps at different time points in response to sham, US (parameter set 1 and 2), and light flashes to both eyes. The US target zone is shown as a black circle. (**B**-**C**) Responses of the targeted region of visual cortex to sham, different US parameters (parameter set 1 [B] and parameter set 2 [C]), and light flashes (n=4 animals, one-way ANOVA ****p < 0.0001, Tukey’s post comparison). Mean trace is solid and SEM is shaded. Bar graph values represent mean ± SEM.

## DISCUSSION

This study presents a new conditionally deafened mouse model for studying the effects of FUS on neural activity in vivo and detailed characterization of the cortex-wide calcium dynamics of deafened mice in response to continuous FUS with 500 ms PD, at pressures ranging from 100 kPa to 2400 kPa. In clear skull preparation, by eliminating the auditory confounds, we observed a significant reduction in off-target brain activation with 270 kHz FUS at pressures up to 500 kPa, and complete elimination of off-target activation with 916 kHz FUS at pressures up to 900 kPa. However, we did observe residual brain activity for the 270 kHz frequency at high pressures of 500 and 900 kPa, which may be attributed to direct activation of somatosensory organs due to the wide beam of the 270 kHz FUS.

Using the TPX window preparation with 916 kHz continuous FUS for 500 ms PD, we observed pressure-dependent off-target and/or localized effects (**Figure 8A**). After eliminating the auditory confounds, we observed localized temperature-dependent fluorescence dimming starting from pressure at 400 kPa due to fluorophore heating (**Figure 4**), which is consistent with a previous report.^33^ When we increased the pressure to 1000 and 1300 kPa, we occasionally observed bilateral off-target brain activation in conjunction with dimming of ipsilateral fluorescence (**Figure 5**). When we further increased the pressure to 1600 kPa or above, we observed strong ipsilateral disruptive depolarization accompanied by contralateral off-target sensory activation (**Figure 6**). To understand the off-target residual sensory effects in deafened mice induced by 916 kHz FUS (**Figures 2, 5, and 6A**), we created a finite element model (FEM) of a mouse, and used it to investigate the response of mouse models to the transcranial UNM of the visual cortex (**Figure 8B**). We found that the induced wave pattern is complex and has both localized and delocalized components, suggesting that peripheral skin receptors and photoreceptors may experience displacements and stresses at levels that could result in the activation of ascending somatosensory and visual pathways, as reported in some peripheral UNM studies.^34,39,41,45^ We also computed the off-target stresses at a small region on the mouse neck, where we observe that the shear stresses (i.e., von Mises stresses), unlike pressures, exhibit a strong frequency dependence in their response (**Figure 8C and 8D**).^46^

**Figure 8.**
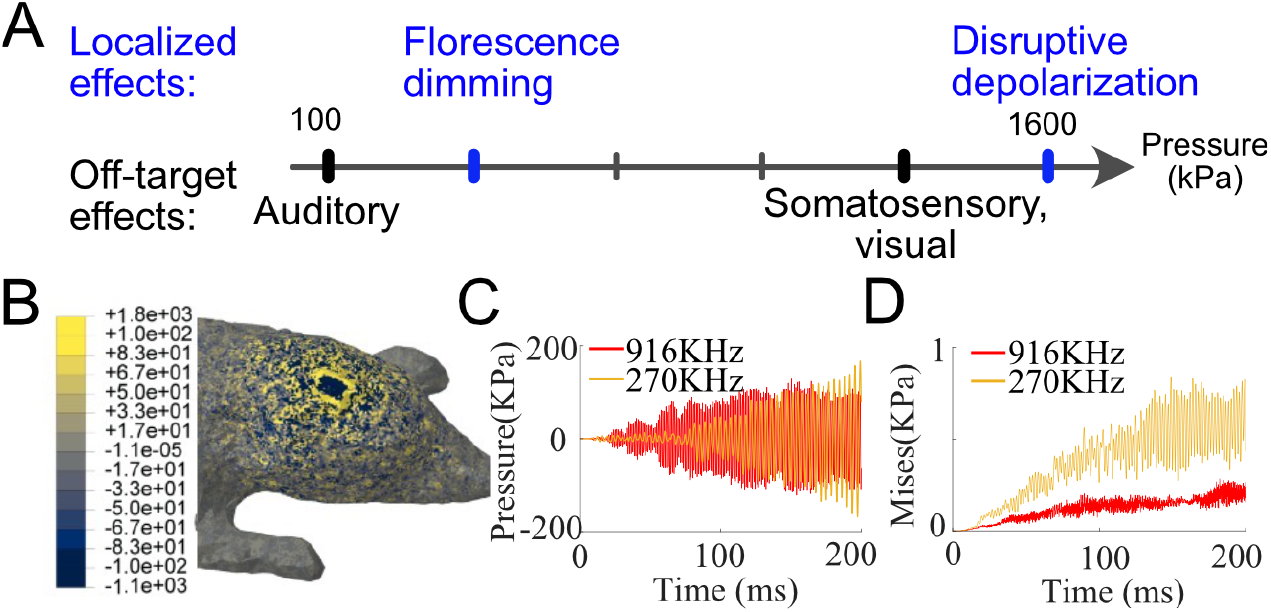
Localized and non-localized effects and potential mechanisms of in vivo UNM in mice. (**A**) US can elicit auditory confounds at low pressure. In deafened mice, the auditory confounds can be greatly reduced or eliminated. However, the thermal confounds at the focus, other sensory confounds, and destructive depolarization will be observed as the pressure of the US increases. (**B**) Result of finite element method (FEM) model of a mouse, where a FUS with peak amplitude of 1000 kPa and frequency of 916 kHz is applied. The results depict the pressure contours after 200 microseconds of explicit dynamics simulations. Note the pressure is widely distributed on the head. (**C**-**D**) Time history of off-target pressures and von Mises stress at a region on the face, obtained from two separate FEM simulations with identical FUS peak amplitudes of 1000 kPa but distinct frequencies.

We found no evidence that FUS evokes direct calcium responses in the live brain (**Figures 3 and 4**) at parameters that were able to do so in cultured neurons (**Figure S6**).^31^ This discrepancy may be due to biophysical differences, as the brain may experience different mechanical and acoustic conditions than cultured neurons. Moreover, the expression levels of the US-sensitive and amplifier channels identified in cultured embryonic neurons (TRPP1/2, TRPC1, and TRPM4) may be different in the adult mouse brain. Future studies may explore cell-type responses to FUS in fully deafened animals to better understand this discrepancy.

Our results do not conclusively demonstrate that FUS is unable to produce direct brain activation. For example, although a comprehensive set of continuous FUS parameters had been tested, we may have overlooked certain parameters, such as even higher center frequency (> 1 MHz) and shorter PDs (< 500 ms). Furthermore, moderate pressure FUS may have generated weak activation masked by temperature-dependent fluorescence reduction. Similarly, high pressure FUS may have produced brain activation that was overwhelmed by the intense seizure-like depolarization attributed to cellular damage. Finally, there might be subcortical activation and finer cortical activation that cannot be observed by wide-field calcium imaging.^21,47^ Therefore, other neural recording approaches, such as functional US imaging or two-photon microscopy, may be needed in future UNM studies to record FUS-elicited neural activity in deep brain or at single neuron resolution.^19,48^

Our study employed a conditionally deafened animal model, which could be useful for sonogenetics studies, which have reported localized brain activation with FUS by expressing or overexpressing various mechanosensitive and/or thermosensitive ion channels (MscL, Prestin, TRPA1, TRPV1) in the brain of normal hearing mice.^36,48–51^ However, it is crucial to use models like our conditionally deafened mice because our previous and current experimental and modeling results have shown that transducers at center frequencies lower or around 1 MHz could elicit strong auditory confounds and off-target brain activation.^20,21,46^ Moreover, researchers should also pay attention to somatosensory and visual confounds, as high-pressure FUS may still activate ascending non-auditory pathways in deafened animals (**Figures 5 and 6**). To distinguish brain activation from disruptive depolarization, it is necessary to perform histology and confirm that the FUS parameters leading to brain activation do not cause any brain damage. In conclusion, UNM holds great potential in treating various brain disorders,^52,53^ and it is always advisable to stay cautious by imaging whole cortical or brain activity and performing control experiments to ensure that the observed brain activation is localized and not confounded.

## METHODS

### Experimental model and subject details

For this study, mice were used in accordance with animal procedures approved by the Institutional Animal Care and Use Committee at the California Institute of Technology. For control experiments, transgenic mice, C57BL/6J-Tg(Thy1-GCaMP6s)GP4.12Dkim/J (The Jackson Laboratory, Stock No. 025776) were used. Strain B6.Cg-Pou4f3tm1.1(HBEGF)Jsto/RubelJ (The Jackson Laboratory, Stock No. 028673) and C57BL/6J-Tg(Thy1-GCaMP6s)GP4.12Dkim/J were crossed to breed the double transgenic heterozygous Pou4f3^+/DTR^ x hemizygous Thy1-GCaMP6s mice, which were used for the deafening procedures.

### Animal surgery

Anesthesia was induced by placing mice in a clean induction chamber and delivering 5% isoflurane. The animal’s hair was removed and was then placed in a stereotax. The head was held steady using ear bars and a nose cone. Anesthesia was maintained via delivery of isoflurane (1.5%-2%) in the nose cone. The body temperature was maintained using a heating pad. Both eyes were protected using ophthalamic ointment. The residual hair was removed using hair removal cream and exposed scalp sterilized using chlorhexidine. Local anesthetic bupivacaine 0.5% was injected just under the scalp and left for 1 minute before any incisions. The skull was then exposed via an incision along the midline and laterally above the cerebellum.^21^ For clear skull preparation, a thin layer of super glue was evenly applied to the exposed skull.

To avoid the aberrance from the skull, in some experiments, we replaced bilateral partial skull with an acoustically and optically transparent polymethylpentene plastic (TPX, Mitsui Chemicals, Minato City, Japan), 0.125 mm in thickness. For TPX implantation surgery, the animals were given subcutaneous injections of an osmotic diuretic

Manitol 20% (≤ 0.5ml) to prevent brain swelling. The size of the TPX window on each side is approximately 7 mm × 4 mm (−4 to +3 mm along the antero-posterior axis; +1 to +5 mm from the midline along the lateral axis) to expose major motor, somatosensory and visual cortices. To prevent heating of cortex, the drilling of the craniotomy was done slowly, sterile saline rinse between short periods of drilling. Angled forceps were used to remove the skull pieces carefully. Next, a pre-cut TPX window was positioned on top of the exposed brain and sealed with tissue adhesive (Vetbond, veterinary grade, 3M).

### Administration of diphtheria toxin

Diphtheria toxin was used in accordance with the procedures approved by the Institutional Biosafety Committee at the California Institute of Technology. We used the same protocol described in Golub’s study.^29^ Adult mice received two intramuscular injections of diphtheria toxin (Sigma-Aldrich D0564-1MG) at 50 ng/g, spaced 2 days apart. Mice received 0.4 ml of lactated Ringer’s solution by subcutaneous injection once or twice daily on days 3–6 after the first DT injection. Between days 1 and 6 after the first DT injection, moist food was provided and was supplemented with high-calorie gel (Tomlyn/Vétoquinol from Nutri-Cal).

### In vivo experimental preparation

Each experiment day, anesthesia was induced by placing mice in a clean induction chamber and delivering 5% isoflurane. As soon as voluntary movement ceased, mice were quickly moved to the UNM setup and maintained at 1-2% isoflurane for preparation. The headplate of the animal was fixed to two aluminum bars. The body of the animal was restrained to a platinum plate with a piece of tape. The skull was thoroughly rinsed and cleaned with sterile saline. The ultrasound transducer was immersed into a 3-D printed cone filled with degassed water. The cone, angled 45 degrees from parallel, was brought to the approximate target (i.e., left or right visual cortex) with a manual 3-axis micrometer (XYZ). Degassed ultrasound gel was used to couple the cone and the brain. The gel was then flatten with a glass plate for optical access. Bilateral copper bars were inserted into the gel mound to maintain the gel temperature at 37°C. The other end of each copper bar was sealed in a 3-D printed tube filled with circulating warm water. Before acquiring images, the isoflurane was turned off and the mouth cone was retracted to allow the mice breathing fresh air. Experiments started after the animals were fully awake and had voluntary movements.

### Experimental protocol design

All imaging animals underwent 50 blocks of experiments. In each block, a trial of each stimulus was presented once in random order. The ISI was 20 s for all experiments except 150 s in Figure 5. In Figure 1, the stimuli in each block were sham, light flash to both eyes, and audible broadband noise to both ears. In Figures 2, 3, 4, and 7, the stimuli in each block were sham, ultrasound at different pressures, and light flash to both eyes. For Figure 5, due to the prolonged time, each block only had one stimulus (sham, US at 1000 kPa, US at 1300 kPa). In Figure 5, due to the long ISI, each block included only one stimulus, which could be either sham, US at 1000 kPa, or US at 1300 kPa. In Figure 6, only one or five trials were presented. In Figures 2 and S4B, the transducer was operated at its fundamental frequency of 270 kHz, while in all other experiments it was operated at 916 kHz. Continuous ultrasound with a PD of 500 ms was utilized for all experiments, except for Figure 7 and S4B, in which we examined some pulsed ultrasound parameters.

### Experimental control

Experiments were controlled by custom software, written in LabVIEW (National Instruments, Austin, TX) and MATLAB (Mathworks, Natick, MA). A PXIe chassis (PXIe-1073) and a function generator (FxnGen) (PXI-5421), both from National Instruments, were used to generate a ramped broadband noise waveform which was amplified by a power amplifier (SA1, Tucker-Davis Technologies, Alachua, FL) to drive an open-field magnetic speaker (MF1, Tucker-Davis Technologies, Alachua, FL). The speaker-ear system was calibrated using a condenser microphone system (PS9200KIT, ACO Pacific, Belmont, CA).^20^ The light flashes were generated by two white LEDs driven by an Arduino UNO. Another Arduino (Mega 2560) was used to trigger the function generator of audible sounds, Arduino UNO of LEDs, wide-field camera, and the signal generator of ultrasound. This Arduino communicated with Matlab in a PC, which randomized the stimuli in each block, to trigger the appropriate stimulus (sham, sound, light, or ultrasound) accordingly.

### Data acquisition and analysis

The construction of the wide-field calcium imaging system has been described in the previous study.^21^ In the present study, images were collected at 20 Hz using a camera (pco.panda 4.2, PCO, Kelheim, Germany) running on the external trigger mode from an Arduino. The exposure of each image was set to 39 ms. One hundred frames (i.e., 5 second recording) were acquired before the on-set of stimuli, where the first and last frames were abandoned when calculating the baseline fluorescence F0 to avoid shutter artifact and stimulus onset noise. After that, 200 or 2900 frames (i.e., 10 or 145 s recording) were acquired for recording evoked calcium responses to different types of stimuli. Each image frame was spatially filtered with a 20 pixel (∼500 μm) square filter to reduce noise. Relative fluorescence intensity changes were calculated as dF/F = (F-F0)/ F0. For clear skull experiments (**Figures 1-3, S1-2**), wide-field images were transformed to Common Coordinate Framework atlas coordinates (CCF, v3 ©2015 Allen Institute for Brain Science) by affine transformation based on manually selected control points.^54^ Cortical areas such as visual, auditory, somatosensory and motor cortices were directly used as ROIs. CCF remapping was not performed for the TPX window experiments (**Figures 4-7**) as the control points were not visible. Hence, the ROI for FUS focus was manually selected based on the negative fluorescence region where FUS focal heating occurred. The recovery time (**Figure 5**) was defined as the time for the fluorescence signal and temperature return to the mean ± 3 × SD of baseline.

### In vitro experimental preparation

The detailed method of primary neuron preparation has been described in the previous study from our laboratory.^31^ The cortical neurons collected from embryonic day 18 C57BL/6J mice (The Jackson Laboratory) were seeded on the top of poly-d-lysine (Millipore Sigma, 70-150k mol wt) pre-coated Mylar dish at a density of 300 cells/mm^2^, and maintained in Neurobasal medium (Thermo Fisher Scientific) supplemented with B27 (2% v/v, Thermo Fisher Scientific), GlutaMax (2 mM, Gibco), glutamate (12.5 μM, Sigma) and penicillin/streptomycin (1% v/v, Corning) in a humidified incubator with 5% CO2 and 37 °C. Half of the medium was changed with the fresh medium without glutamate every 3 days, and neurons were used for ultrasound stimulation experiments after 11–12 days from the seeding. Water bath temperature was maintained to 37 °C during the US stimulation experiment. For calcium imaging, Syn-driven GCaMP6f as a calcium sensor was delivered to neurons via AAV1 viral vector transfection (Addgene 100837-AAV1, 1E10 vp/ dish) at 4 days in vitro. For the experiments characterizing the temperature dependence of fluorescence in vitro, a HEK293 cell line constitutively expressing GCaMP6f under the CMV promoter were cultured in Dulbecco’s modified Eagle’s medium (DMEM) supplemented with 10% tetracycline-free fetal bovine serum (FBS) and penicillin/streptomycin. Cells were trypsinized, resuspended in Opti-MEM (Thermo Fisher Scientific), and added to 200μL qPCR tubes. The qPCR machine was programmed to perform a temperature ramp from 37 °C to 42 °C in 1-degree increments, while the GCaMP fluorescence was read-out simultaneously using the corresponding FAM fluorescence filters.

### Ultrasound generation and calibration

Details of ultrasound generation and calibration are described in our previous study.^20,21^ A single element transducer (H-115, Sonic Concepts, Bothell, WA) was used in all experiments. The transducer can run at its first harmonic (270 kHz) and third harmonic (916 kHz) with different matching boxes, which were driven by a 200 W RF amplifier (E&I 2200, Electronics & Innovation, Rochester, NY). The amplifier was triggered by a waveform generator (33500B series, Keysight Technologies, Santa Rosa, CA).^20^ Calibration was done in a large water tank with a fiber-optic hydrophone system (FOH, Precision Acoustics, Dorchester, UK). Unless stated otherwise, pressures listed in the manuscript refer to underated peak negative pressure measured in a water tank.

### Finite element modeling

We adopt the geometry for our three dimensional finite element model from the co-registered x-ray CT of a male mouse developed in the Digimouse project.^46,55^ This model contains all the major anatomical parts of the mouse body: skeleton, whole brain, heart, lungs, liver, stomach, spleen, pancreas, kidneys, testes, bladder, muscles and skin. We have discretized the mouse anatomy into a finite element model with more than 10 million three-dimensional linear tetrahedral elements. We have also adopted organ-dependent material properties including elasticity and viscoelasticity. We subject a region on the skin on top of the skull to harmonic pressure with intensity of 1 MPa. In order to have the US pressure focused inside the brain, the pressure is imposed in a phased array manner, where the phase lag is introduced in a radial direction. We then solved the initial boundary-value problem of small-strain elasticity with recourse to the explicit finite element method (Abaqus/Explicit, Dassault Systèmes Simulia, France).

### Quantification and statistical analysis

All raw data was processed using custom code written in MATLAB. Statistical analysis was performed using the program GraphPad Prism 9 (GraphPad Software, San Diego, CA). All the figures were plotted using Matlab and Prism. Maximum dF/F signal was used for group comparisons.

## ACKNOWLEDGEMENTS

The authors thank Joseph Wekselblatt, David Mittelstein, and Pierina Barturen-Larrea for helpful discussion, and members of the Shapiro and Tsao labs for assistance with experiments. This research was supported by NIH BRAIN Initiative grant R24MH106107 (Co-PIs D.Y.T. and M.G.S.), RF1MH117080 (to M.G.S.), NIH grant DP1 EB033154 (to M.G.S.), and the Howard Hughes Medical Institute (D.Y.T.). Related research in the Shapiro Laboratory is also supported by the Howard Hughes Medical Institute and the Packard Fellowship in Science and Engineering.

## AUTHOR CONTRIBUTIONS

H.G., D.Y.T., and M.G.S. conceived the study. H.G and M.G.S. designed all experiments and H.G performed all the in vivo experiments and analyzed the data. D.W. and S.J.Y. performed the in vitro experiments in cell cultures and analyzed the data. H.S. performed the modeling. T.S. and M.G.S. initialized the idea of using the double transgenic mouse model. H.G. and M.G.S. wrote the manuscript with input from all authors.

## COMPETING INTERESTS

The authors declare no competing interests.

## DATA AND MATERIALS AVAILABILITY

Raw data are available upon reasonable request to the authors.

## Notes

### Competing Interest Statement

The authors have declared no competing interest.

