## Supplementary information for "Effects of focused ultrasound in a “clean” mouse model of ultrasonic neuromodulation"

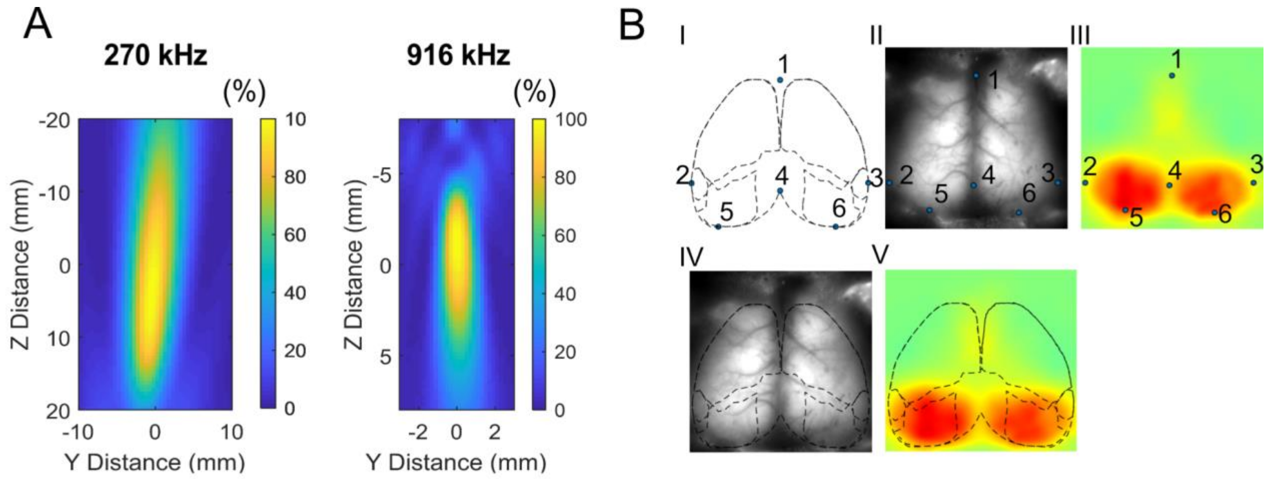


**Supplementary figure 1. Transducer calibration and brain remapping using** Allen Mouse Brain Common Coordinate Framework 3.0. (**A**) Calibration of the transducer (Sonic Concepts, model H-115) at its fundamental frequency 270 kHz and the third harmonic 916 kHz, which have a lateral full width at half maximum [FWHM] for pressure of 8.85 mm and 1.4 mm, respectively. (**B**) The brain remapping used six control points (I) in the top view of Allen Mouse Brain CCF and another six control points from the raw wide-field image and the calcium activity map (II an dIII), where the points can be manually adjusted. After remapping (IV-V), different cortical regions can be identified and the regional responses (visual, auditory, somatosensory and motor) can be calculated using the remapped ROIs.

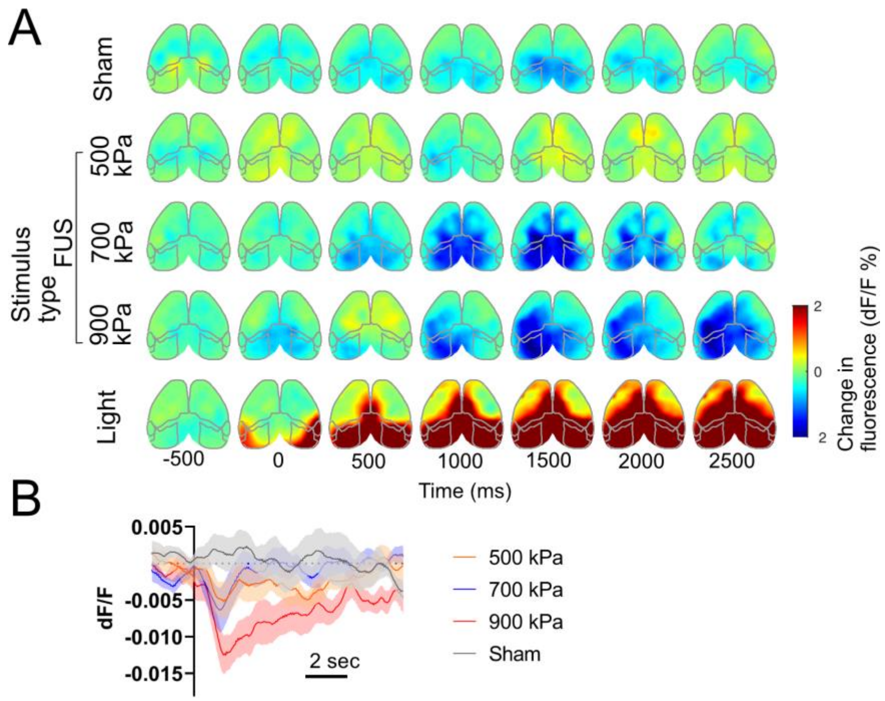


**Supplementary figure 2. Ultrasound reduces calcium response at the focus in deafened mice with clear skull preparation**. (**A**) A representative example of cortical activation map at different time points in responses to sham, ultrasound (500, 700, 900 kPa; 500 ms PD), and light flashes to both eyes. The ultrasound was delivered through the skull in the clear skull preparation. (**B**) Responses of the focus to different ultrasound parameters and sham (n=4 animals). Clear negative and long-lasting fluorescence signals can be observed for pressures of 700 kPa and 900 kPa.

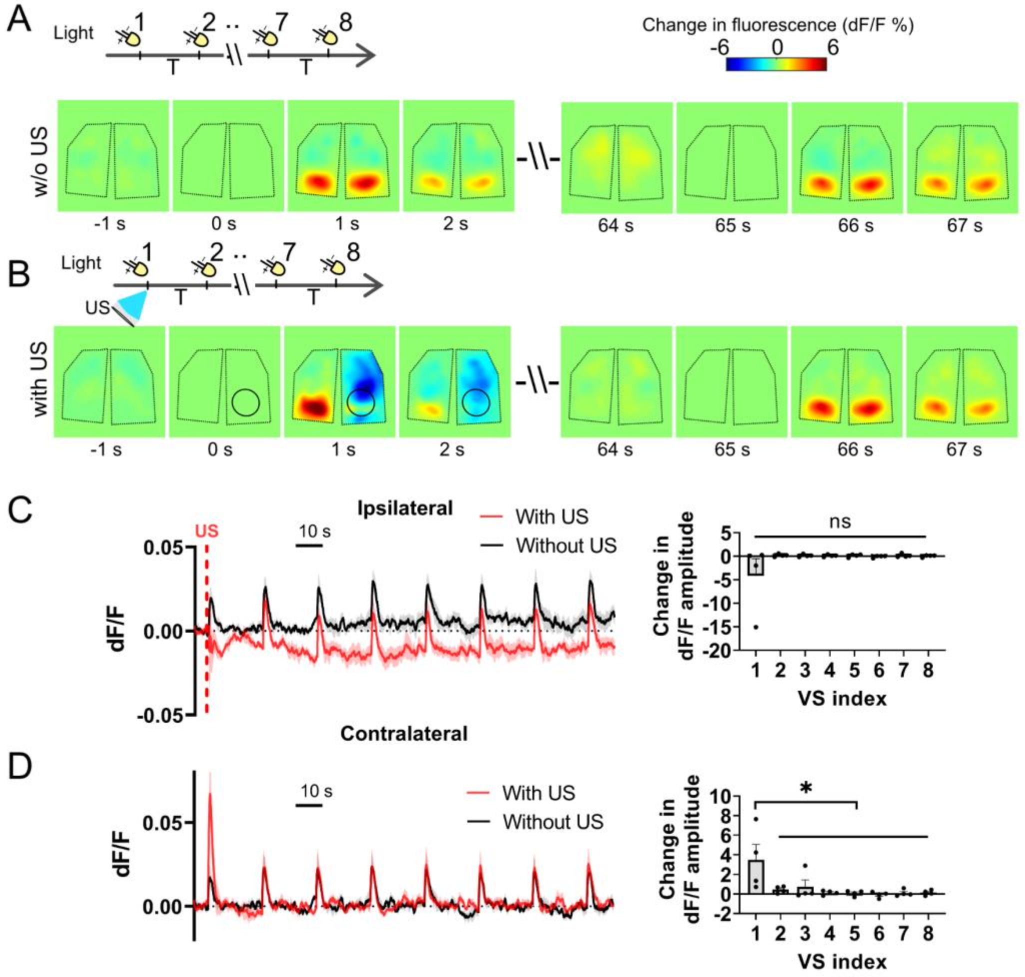


**Supplementary figure 3. Paired ultrasound and light flashing stimulation enhance bilateral and reduce ipsilateral visual evoked calcium responses.** (**A**) Diagram of the light flashing (80 ms duration) stimulation, where eight consecutive light flashes were presented at an interval of 20 s (light-only). A representative cortical activation map is shown. (**B**) Diagram of the paired ultrasound and light flashing stimulation, where eight consecutive light flashes were presented at an interval of 20 s and a pulse of ultrasound (916 kHz, 1300 kPa, 500 ms PD) was delivered simultaneously with the first light flash (light-US). A representative cortical activation map in response to the first and fourth light flash in the two diagrams is shown. The ultrasound target zone is shown as a black circle. The black polygons represent the boundaries of the TPX windows. (**C** and **D**) Response time course to light flashes (with or without ultrasound) at the ipsilateral focus and contralateral counterpart (n =4 animals, 50 blocks for each animal, each block consists of one light flash trial and one paired stimulation trial). The change in peak dF/F of the eight visual responses was normalized to that of the light-only paradigm, respectively. The contralateral visual response was significantly enhanced ($p < 0.05$) following the FUS stimulation while returning to normal from the second visual response. In C, FUS only abolished the first ipsilateral VER and did not change the relative peak amplitudes of the following seven ipsilateral VERs. However, it resulted in a negative shift of the baseline of focal calcium activity, indicating that some neuronal inhibition may exist.

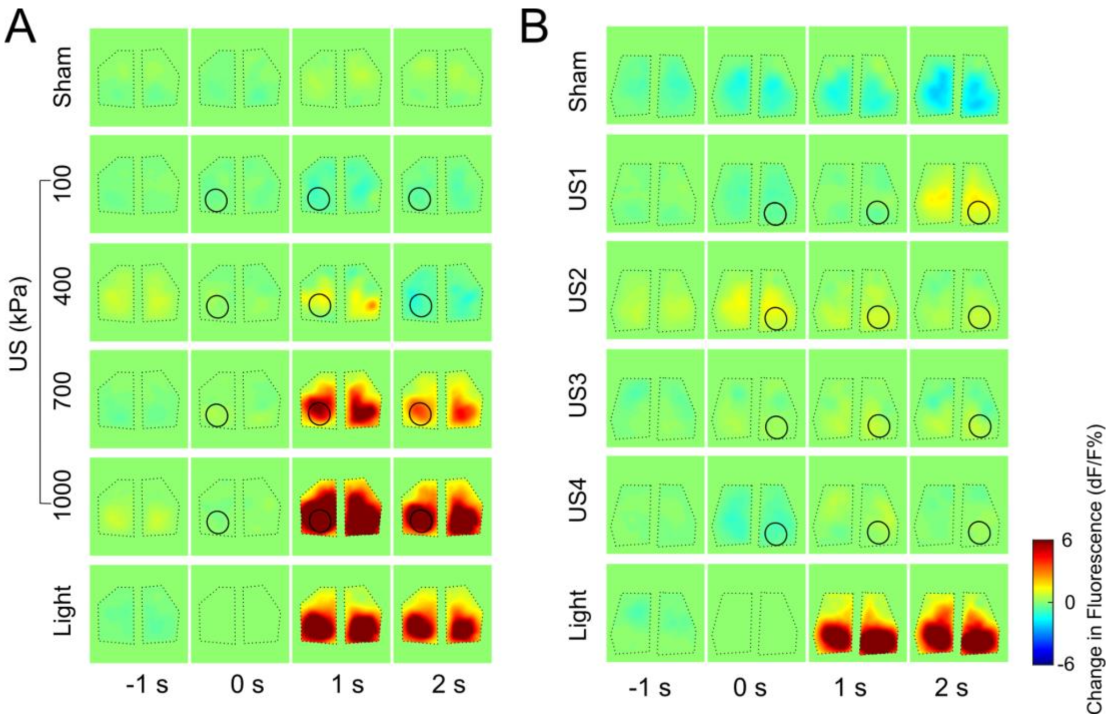


**Supplementary figure 4. FUS elicits auditory confounds in normal hearing TPX mice but not deafened TPX mice.** (**A**) A representative cortical map of extensive cortex wide brain activation in the normal hearing mouse through the TPX window in response to continuous ultrasound (916 kHz, 500 ms, pressure at 100, 400, 700, and 1000 kPa). The focus was on the left visual cortex. (**B**) A representative cortical map of eliminated off-target brain activation to pulsed ultrasound (270 kHz, Parameter set 1 in Table 1) in the deafened mouse. The focus was on the right visual cortex.

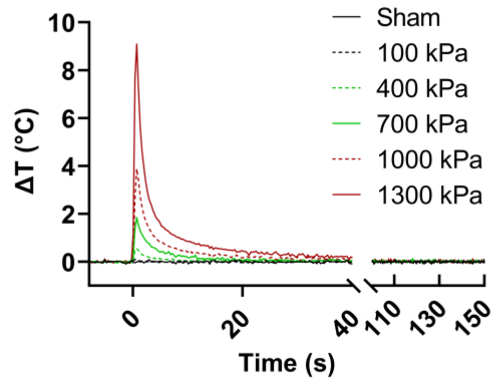


**Supplementary figure 5. Temperature increases at the brain surface under the TPX window during UNM.** The PD was 500 ms for all the pressures. Peak temperature increases were 0.10, 0.56, 1.84, 3.89, and 9.09 °C for 100, 400, 700, 1000, and 1300 kPa, respectively, which were greater than the mean plus three times the standard deviation of the sham. Pressure of 100 kPa did not increase the temperature significantly. Each temperature curve was averaged across 10 trials.

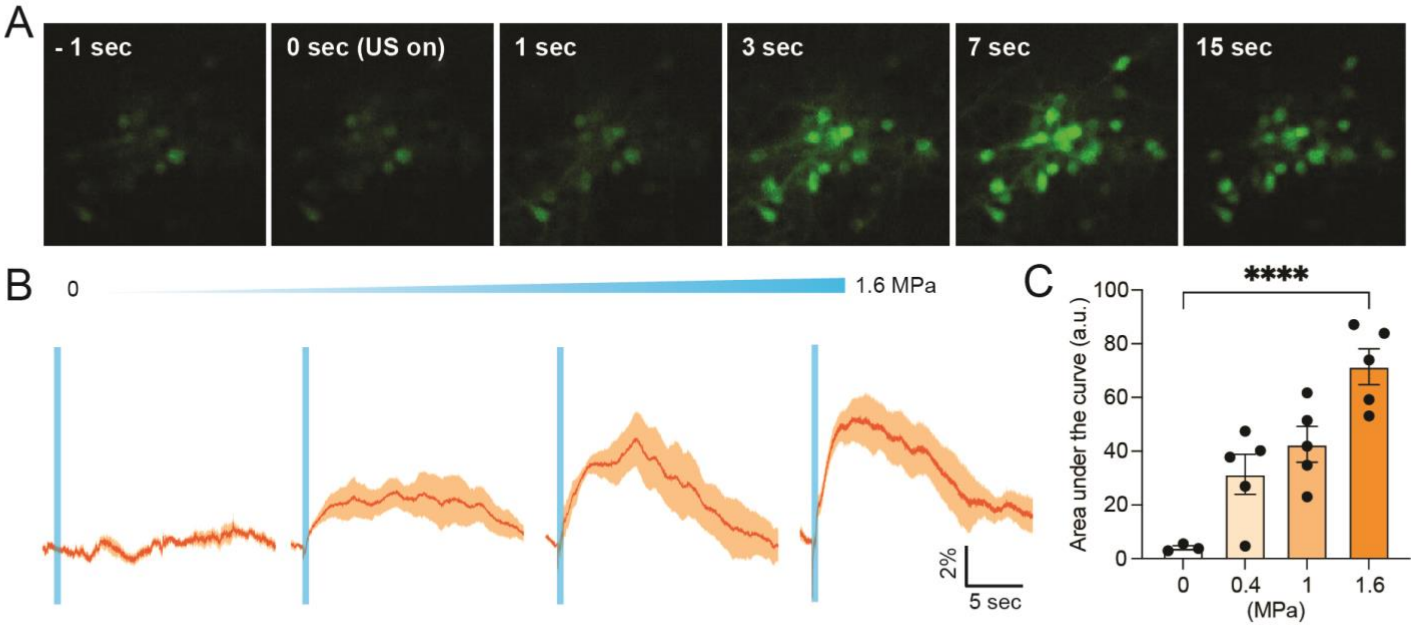


**Supplementary figure 6. 916 KHz ultrasound stimulation to the cultured neurons.** (**A**) Time lapse calcium images of GCaMP6f neurons before and after stimulation. (**B**) Calcium responses (mean trace is solid, and SEM is shaded.) and (**C**) quantification of the response as function of ultrasound intensity (mean ± SEM, n = 3 (0 MPa), 5 (0.4-1.6 MPa) independent experiments each, Tukey's multiple comparison test after one-way ANOVA, $p < 0.0001$).
